# Spatial reorganization of *Escherichia coli* chromosome contextualizes triclosan stress-related genetic, epigenetic and transcriptome changes

**DOI:** 10.1101/2023.07.11.548559

**Authors:** Dipannita Ghosh, Benjamin A. Evans, Perumal Vivekanandan

## Abstract

Changes in the spatial organization of bacterial chromosomes under stress conditions and its biological implications remain poorly understood. We mapped the structural landscape of wild-type and Δ*dcm E. coli* chromosomes under triclosan stress using Hi-C to identify triclosan-induced chromosomal interaction domains (CIDs). Two CIDs were common to the wild-type and Δ*dcm E. coli*, including a CID with a common boundary at *fabI* gene, which encodes the triclosan target. All mutations and structural variants under triclosan stress were observed within or in close proximity to triclosan-induced CIDs. Absence of Dcm methylation impacts both short- and long-range interactions in triclosan stress. Single-base resolution methylome maps reveal hypermethylation of adenines (in wild-type and Δ*dcm*) and cytosines (in wild-type) in the two common triclosan-induced CIDs. Furthermore, global gene expression profiling identified enrichment of highly expressed genes within the two common CIDs. Our findings suggest that stress-induced CIDs in *E. coli* are hotspots for genetic variations and are associated with enhanced transcriptional activity and hypermethylation of Dam/Dcm motifs.

## BACKGROUND

Ubiquitous use of biocides, such as triclosan, and subsequent bacterial resistance to them has led to serious concerns regarding their role in the development of cross-resistance to clinical antimicrobials in bacteria (1–5). In recent years, there is an increasing interest to understand non-classical mechanisms which can confer survival advantage to bacteria in the presence of antimicrobial agents, such as epigenetic modifications (6). A recent study which investigated the role of Dam methyltransferase in the development of triclosan resistance has shown that Dam-mediated methylation can contribute to cross-resistance to aminoglycosides by altering the expression of the *acrAD-tolC* efflux pump (4). Bacterial DNA also gets methylated at cytosine residues by cytosine methyltransferases, of which the most notable is Dcm. Dcm-mediated methylation has also been shown to regulate *sugE* expression (7), which influences bacterial resistance to quaternary ammonium compounds and ethidium bromide (7,8). Besides epigenetic modifications, expression of genes can also be regulated by changes in the spatial organization of chromosomes. This is especially well-established in eukaryotic systems (9). In bacterial systems, chromosomes are organized into compact nucleoid structures with associated proteins, which can regulate gene transcription (10). Folding of the DNA inside the cells can facilitate interaction between regions which can be more than 100 kilobases apart, and these interacting regions form insulated structures called chromosomal interaction domains (CIDs) (11–13). CIDs can range from a few kilobases to hundreds of kilobases, and they are usually flanked by highly expressed genes at their boundaries (12–15). External stressors can significantly alter transcription of bacterial genes through changes in DNA superhelicity (16). Environmental changes also elicit nucleoid remodeling through nucleoid-associated proteins (NAPs) such as histone-like protein HU, which is correlated with modulation of bacterial gene expression (17).

Triclosan is a non-antibiotic, anti-microbial (NAAM) biocide which is widely used in personal products. In this study, we investigated the effect of triclosan-induced stress on the chromosome structure of *E. coli* and how these topological changes are linked with genome-wide methylome landscape as well as the transcriptome of the cells under active stress conditions. We adapted *E. coli* wild-type and Δ*dcm* strains to increasing concentrations of triclosan and probed the phenotypic changes in the adapted strains. This was followed by investigation of the chromosomal reorganization brought about by triclosan stress conditions in the adapted strains. We observed that triclosan stress led to the formation of distinct CIDs in *E. coli* which are hotspots for triclosan-induced mutations as well as structural variations. We also found that both adenine (GA^m^TC) and cytosine (CC^m^WGG) methylome landscape changes caused by triclosan stress mirror the triclosan-induced topological changes. Further interrogation revealed significant upregulation of genes within triclosan stress-induced chromosomal domains compared to that under normal conditions. These results indicate that under stress conditions, bacterial chromosome can form distinct topological structures and these structures are hotspots for genetic variations and enhanced DNA methylation and gene expression.

## RESULTS

### Phenotypic changes associated with adaptation to triclosan in *E. coli* wild-type and Δ*dcm* strains

We assessed the minimum inhibitory concentration (MIC) of triclosan for both *E. coli* wild-type and Δ*dcm* strains, which was found to be 0.5 µg/ml in our laboratory conditions. Both strains were adapted to concentrations of 32 µg/ml. In wild-type *E. coli*, we observe that there is no significant difference in growth between control and triclosan-adapted strains (Supplementary Figure S1A & B). Also, consistent with an earlier report (18), we find that the absence of the Dcm methyltransferase does not directly have any impact on *E. coli* growth under normal conditions. However, the absence of a functional Dcm leads to a modest but significant reduction in the growth rate of triclosan-adapted Δ*dcm* strain compared to control (Supplementary Figure S1B). We did not observe any change in the growth curves for the adapted and control strains under normal conditions (Supplementary Figure S1A), indicating that entry to different phases of bacterial growth is not impacted by adaptation to triclosan. However, in the presence of triclosan stress (at 32 µg/ml), we do observe that both the adapted wild-type and Δ*dcm* strains have a prolonged lag phase, the extent of which is higher in the latter (Supplementary Figure S1C). Environmental stress conditions have been associated with an increased lag phase (19).

Adaptation to biocides is often observed to be associated with cross-resistance to antibiotics in bacteria (1–4). Resistant bacteria are also known to develop collateral sensitivity to unrelated antibiotics, as a result of evolutionary trade-off to facilitate enhanced survival (20). Our antimicrobial susceptibility testing results (Supplementary Figure S1D) are in line with earlier findings showing a similar increase (1,3,4) as well as decrease (21) in susceptibilities to different antibiotics in both triclosan-adapted wild-type and Δ*dcm E. coli*. Interestingly, we found that the decrease in susceptibility to erythromycin, tetracycline and rifampicin after triclosan adaptation is more pronounced in the Δ*dcm E. coli* compared to wild-type. It appears that the absence of Dcm methyltransferase in *E. coli* leads to higher survivability when challenged with antibiotics targeting RNA and protein synthesis.

### Triclosan stress incudes changes in the higher-order chromosome structure of *E. coli*

To investigate if triclosan-induced stress alters the spatial organization of the *E. coli* chromosome, we performed Hi-C analysis on triclosan-adapted wild-type and Δ*dcm* strains (growing in media with 32 µg/ml triclosan) along with their respective mock-adapted controls. The quality assessment of the paired-end reads and Hi-C matrices can be found in Supplementary Table 1.

The contact maps of interaction frequencies for all the *E. coli* strains contain a single prominent diagonal which represents high levels of interaction between adjacent loci in the same chromosomal arm (Figure 1A). The lack of a second diagonal in the maps is consistent with earlier findings (11,22) that the two arms of replicating *E. coli* chromosome occupy different nucleoid regions organized about a transverse axis, and thus inter-arm interactions are unlikely in replicating *E. coli*. Visual inspection of the contact maps showed more pronounced short-range interactions in triclosan-adapted Δ*dcm E. coli* compared to its corresponding control or the wild-type strains. When we plotted the contact frequencies as a function of genomic distance between interacting loci, we observed that the adapted Δ*dcm E. coli* under triclosan stress has higher levels of interactions in the range of ∼10-100 Kb compared to the other strains (Figure 1B). The plot also revealed a steep decline in interaction frequency with increase in genomic distance for Δ*dcm E. coli*, unlike the rest of the strains. Clearly, absence of a functional Dcm methyltransferase and the resulting absence of cytosine methylation influences short-as well as long-range interactions under triclosan stress conditions.

**Figure 1.**
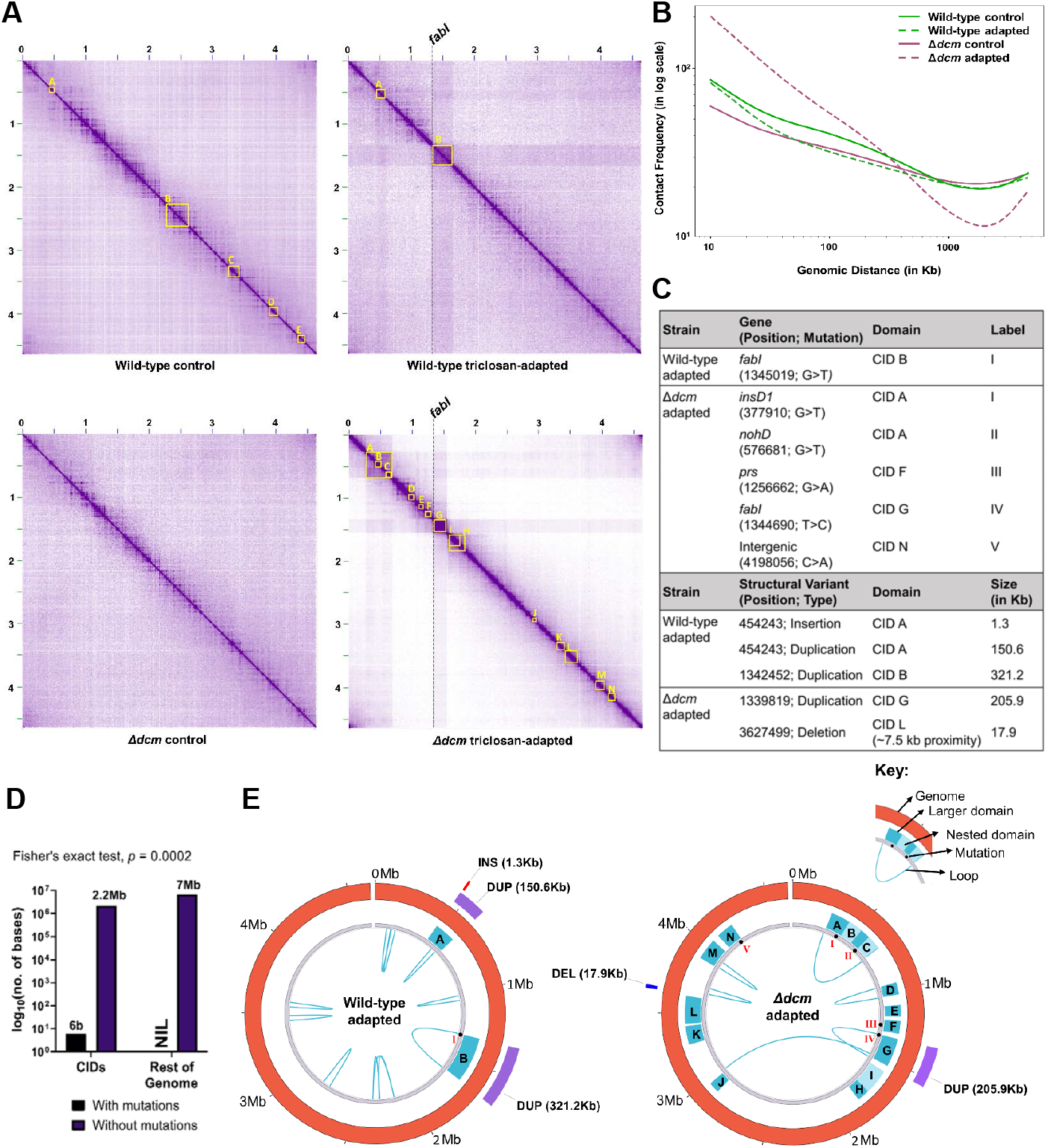
Triclosan stress induced alterations in *E. coli* chromosome topology are hotspots for mutations and structural variations. **(A)** Normalized contact maps of triclosan-adapted wild-type and Δ*dcm E. coli* along with their respective controls. The x- and y-axes correspond to the chromosome coordinates of *E. coli* BW25113. The contact maps are visualized at 10 Kb resolution for whole chromosome and the color intensity is directly proportional to interaction frequency. Chromosomal interaction domains (CIDs) identified at 5 Kb, 10 Kb and 25 Kb resolutions are represented with yellow squares. The CIDs are alphabetically labelled and their exact positions are listed in Supplementary Table 2A. Both triclosan-adapted wild-type and Δ*dcm* strains have at least one CID with boundary at 1.34 Mb locus, which is absent in their respective controls. This 5 Kb locus at 1.34 Mb harbors *fabI* gene. **(B)** Comparison of decay in contact frequency with increase in genomic distance reveals triclosan-adapted Δ*dcm E. coli* has much higher contact between loci separated by ∼10-100 Kb compared to the other three strains and the contact frequency falls sharply as genomic distance between loci approaches ∼1000 Kb. **(C)** Triclosan-adaptation associated mutations and structural variants in wild-type and Δ*dcm E. coli* along with their positions in genome and the associated CIDs. **(D)** Triclosan-adaptation associated mutations are significantly enriched inside chromosomal domains. The bar graphs compare the frequency of triclosan-adaptation associated mutations (both wild-type and Δ*dcm E. coli* combined) within CIDs and the rest of the genome. **(E)** Triclosan-adaptation associated mutations, structural variants, and topological features including CIDs and loops identified from the contact maps are collated into circular plots. All the adaptation-associated mutations and structural variations identified co-localize with CIDs. The start and end loci of these loops can be found in Supplementary Table 2B.

Loci which interact more frequently with each other form chromosomal interaction domains (CIDs). To probe if triclosan stress induces changes in these structural features of the *E. coli* chromosome, we annotated the contact maps with domains identified by Juicer (23) (Arrowhead algorithm) at different resolutions (Figure 1A, Supplementary Table 2A). Five CIDs were identified in the wild-type control strain ranging from 90kb to 360kb, whereas, none were found in the Δ*dcm* control. In the adapted strains under triclosan stress, the wild-type *E. coli* had two CIDs of sizes 145kb and 320kb, while fourteen CIDs were identified in Δ*dcm* strain, with sizes between 75kb to 405kb. We also observed some smaller domains nested inside larger domains specifically in the Δ*dcm* adapted strain, such as CID B and C inside the larger CID A, and CID I present within CID H. Similar nested CIDs have been previously reported in the *Caulobacter* chromosome (12). Interestingly, we found that in both wild-type and Δ*dcm E. coli* under triclosan stress, at least one of the domains identified had a boundary overlapping the *fabI* gene (labelled CID B in wild-type and CID G in Δ*dcm*) at 1.34 Mb locus. FabI or enoyl-acyl carrier protein (ACP) reductase is an essential component of the type II fatty acid biosynthesis pathway and is a target for several antimicrobials, including the biocide triclosan (24). Also, overexpression of *fabI* has been widely reported following adaptation to triclosan (4,25). It is important to note here that while CID B in wild-type and CID G in Δ*dcm* triclosan-adapted *E. coli* have a common boundary containing the *fabI* gene, the other boundaries of the same CIDs are found at different loci in the two strains. This indicates that while the domain boundary at *fabI* is fixed, the other end of the domain can have variable boundaries potentially determined by local transcription profiles and other factors. To assess the transcription profile of genes present at the boundaries of CID B (wild-type) and CID G (Δ*dcm*) in our samples, we performed reverse-transcription quantitative PCR (RT-qPCR) for *fabI, sapD* present at the common locus at 1.34Mb, *clcB, ynfL* for boundary at 1.66Mb in wild-type, and *yddG* and *narU* for boundary at 1.54Mb in Δ*dcm E. coli* (Supplementary Figure S2). The genes at the common boundary at 1.34Mb are significantly upregulated in both wild-type and Δ*dcm* strains under triclosan-stress. However, upregulation of the genes at the other boundaries of the same domains is not significant for all the genes tested. Our findings, showing the *fabI* gene at the common boundary of CIDs present in *E. coli* under triclosan stress, are in agreement with earlier studies that reveal CID boundaries can be characterized by actively transcribed genes among other factors (11,12).

### Triclosan stress associated chromosomal interaction domains are hotspots for genetic variation

We used Snippy v4.6.0 to identify single nucleotide polymorphisms (SNPs) and small insertions/deletions in our adapted and control strains (Figure 1C and Supplementary Table 3A). The control wild-type *E. coli* strain did not have any mutations, while the adapted strain had a nonsynonymous G>T substitution (G93V) in *fabI*, which has been previously reported(4,26). This G93V mutation alters the protein structure resulting in changes in aromatic interactions with triclosan, thereby increasing resistance to triclosan (27). In the control Δ*dcm* strain, we observed three mutations in *dxr* (nt.190993, nonsynonymous, A>T), *selB* (nt.3752869, deletion, CGATCGCGTGG) and *hdfR* (nt.3940599, deletion, T). These mutations, which are unrelated to triclosan adaptation, are also found in the adapted Δ*dcm* strain and are likely due to absence of a functional Dcm methyltransferase. The adapted Δ*dcm* strain harbored four additional nonsynonymous mutations in: (i) an IS2 element transposase, *insD1* (nt.377910, G>T), (ii) a DNA packaging protein NU1 homolog, *nohD* (nt.576681, G>T), (iii) the phosphoribosylpyrophosphate (PRPP) synthetase, *prs* (nt.1256662, G>A), (iv) *fabI* (nt.1344690, T>C or F203L); and an intergenic mutation (nt.4198056, C>A). The *fabI* F203L mutation is known to confer resistance to triclosan by reducing binding affinity for triclosan (28).

Interestingly, we found that all the 6 triclosan-induced mutations (Figure 1C) were seen within CIDs, although CIDs account for only about a fourth of the entire genome (Figure 1D and 1E). Many of the mutations in Δ*dcm* strain appeared to be in close proximity to the CID boundaries, including CID C (576681G>T, *nohD*, ∼3.3 kb), CID G (1344690T>C, *fabI*, ∼4.6 kb) and CID N (4198056C>A, intergenic, ∼16.9 kb). The *fabI* mutation (G93V) in wild-type *E. coli* is also present at the boundary of CID B (∼5Kb). It is important to take into consideration that the highest resolution of these Hi-C maps at which these domains and boundaries were identified is 5kb. Hence, an even higher resolution may help better resolve the true boundaries of these domains. We also mapped the chromatin loops identified by Juicer (HiCCUPS algorithm) to assess if there is any overlap with the triclosan-associated CIDs. We found that in wild-type and Δ*dcm* strains under triclosan-stress, the domains with boundaries harboring *fabI* gene (CID B and CID G respectively), are likely to be formed due to tethering of the boundary loci forming a loop. CID A and CID D in Δ*dcm E. coli* also appear to be loop domains.

Recent studies have revealed that there is a reciprocal relationship between genomic structural variants (SVs) and spatial architecture in eukaryotic systems (29,30). However, their association in prokaryotes remains unclear. In our study, we used long reads from native DNA sequencing (performed in MinION, Oxford Nanopore Technologies) to identify structural variants in triclosan-adapted and control *E. coli* strains (Figure 1C & E, and Supplementary Table 3B). In the adapted wild-type *E. coli*, we found three SVs which were absent in the corresponding unadapted control, including a 1.3 kb insertion at nt.454243, and two duplications at nt.454243 (150.6 kb) and nt.1342452 (321.2 kb). Interestingly, all three of these SVs co-localize with the CIDs associated with triclosan-stress. A similar overlap between triclosan-induced SV and CID is also observed in the Δ*dcm E. coli*, wherein a 205.9 kb duplication at nt.1339819 overlaps with CID G in the triclosan-adapted strain which is missing in the unadapted control. We also found a 17.9 kb deletion at nt.3627499 in very close proximity (∼7.5 kb) to one of the boundaries of CID L (locus 3620000) in adapted Δ*dcm E. coli*. As mobile genetic elements (MGEs) such as Insertion Sequence (IS) elements often act as sources of SV formation (31), we also assessed the presence of MGEs in the SVs identified from the triclosan adapted strains (Supplementary Figure S3 and Supplementary Table 4). In the adapted wild-type *E. coli* strain, we found that the 1.3 kb insertion at nt.454243 has very high sequence identity (93%) with IS421. The 150.6 kb duplication in the same strain at the same locus, nt. 454243, also harbored IS elements IS421, IS3, IS5 and a miniature inverted repeat MITEEc1. It is interesting to note that the 1.3kb SV identified as IS421 element at nt.454243 (locus 450000; starting boundary of CID A) is also present at the end boundary of CID A, at locus 600000. Whether the insertion of IS421 at nt.454243 is a result of close proximity of the two loci forming CID A requires further investigation. While the duplications at nt.1342452 (adapted wild-type) and nt.1339819 (adapted Δ*dcm*) harbor several MGEs, these MGEs are unlikely to be sources of the large-scale duplications as they are not proximal to the boundaries of these SVs. The deletion at nt. 3627499 in the adapted Δ*dcm* strain did not have any MGEs. Since it is known that SVs get introduced at loci with pre-existing contacts (30), it is likely that the SVs observed in our study are formed due to the long-range interactions caused by triclosan stress conditions.

### Hypermethylation of cytosines and adenines within chromosomal interaction domains

DNA methylation and three-dimensional organization of chromosomes are known to be interlinked in eukaryotes, and this may influence downstream processes (32–34). However, in bacteria, the interplay between spatial organization and genome-wide methylome landscape remains poorly understood. In addition, DNA methylation has been associated with several cellular processes in bacteria which can confer survival advantage under adverse conditions(6,7,35), hence, we wanted to investigate the methylome changes associated with adaptation to triclosan stress. Long-reads generated from native DNA sequencing were used to perform modified basecalling with Tombo v.1.5.1 (36) to identify methylation levels of GATC and CCWGG motifs in the *E. coli* genomes.

We assessed the distribution of fraction of modified reads (ModFrac) values for the internal cytosine of CCWGG motifs (Dcm methylation site) in all four strains using kernel density estimation (KDE) in R (Figure 2A). We found that the cytosine ModFrac values for Δ*dcm* strains peak at ∼0.1, whereas for the wild-type strains (control and adapted) which have functional Dcm methyltransferase, the peaks can be seen at ∼0.9. Peaks at 0.1 to 0.15 have been previously documented for unmodified bases, indicating background associated with the modified basecalling toolkit, Tombo (37); these peaks are not considered for further analysis. We analyzed the ModFrac values for the wild-type strains which have functional Dcm methyltransferase. The median modified fraction value of the internal cytosine of CCWGG motifs was marginally higher in the triclosan-adapted wild-type *E. coli* compared to that in control *E. coli* (0.9130 vs 0.9091; Figure 2A). When we probed the expression levels of Dcm methyltransferase in the adapted wild-type *E. coli* under triclosan stress compared to the unadapted control, we observed a ∼2.5-fold increase in *dcm* expression (Figure 2B). Since the difference between the median ModFrac values for adapted and control wild-type strains is very minimal (0.0039), higher expression of *dcm* is unlikely to alter the global CCWGG methylome by itself under triclosan stress. To understand if the chromosomal interaction domains impact Dcm methylation landscape, we plotted a three-color heatmap for the adapted and control strains, where each row corresponds to a methylated cytosine position (inside CC^m^WGG motif) on the chromosome and the color scale represents its respective ModFrac value (Figure 2C). Strikingly, we observed hypermethylation of cytosines in the Dcm methylation sites within the chromosomal domains (CID A and CID B) in the adapted wild-type strain under triclosan stress compared to its control. The median ModFrac values for the Dcm methylation sites within CID A and CID B were significantly higher for the adapted strain compared to the Dcm sites found in the corresponding regions of the unadapted control (Figure 2C extreme right summary plots). Also, median ModFrac values for the Dcm sites in both CID A and B are significantly higher than that for the rest of the genome of the adapted wild-type *E. coli* under triclosan stress; this was not observed for the unadapted control (Supplementary Figure S4). Furthermore, ModFrac values for the Dcm motifs mapping to the genomic regions outside the domains (i.e., CID A and CID B), were comparable between the adapted wild-type strain and its corresponding control (Figure 2D). Together, our results show that higher levels of Dcm-mediated cytosine methylation in the adapted wild-type *E. coli* are exclusive to the triclosan-induced chromosomal domains (CID A and CID B). These findings suggest that triclosan stress-induced CIDs in the *E. coli* chromosome are amenable to hypermethylation of cytosines in Dcm motifs.

**Figure 2.**
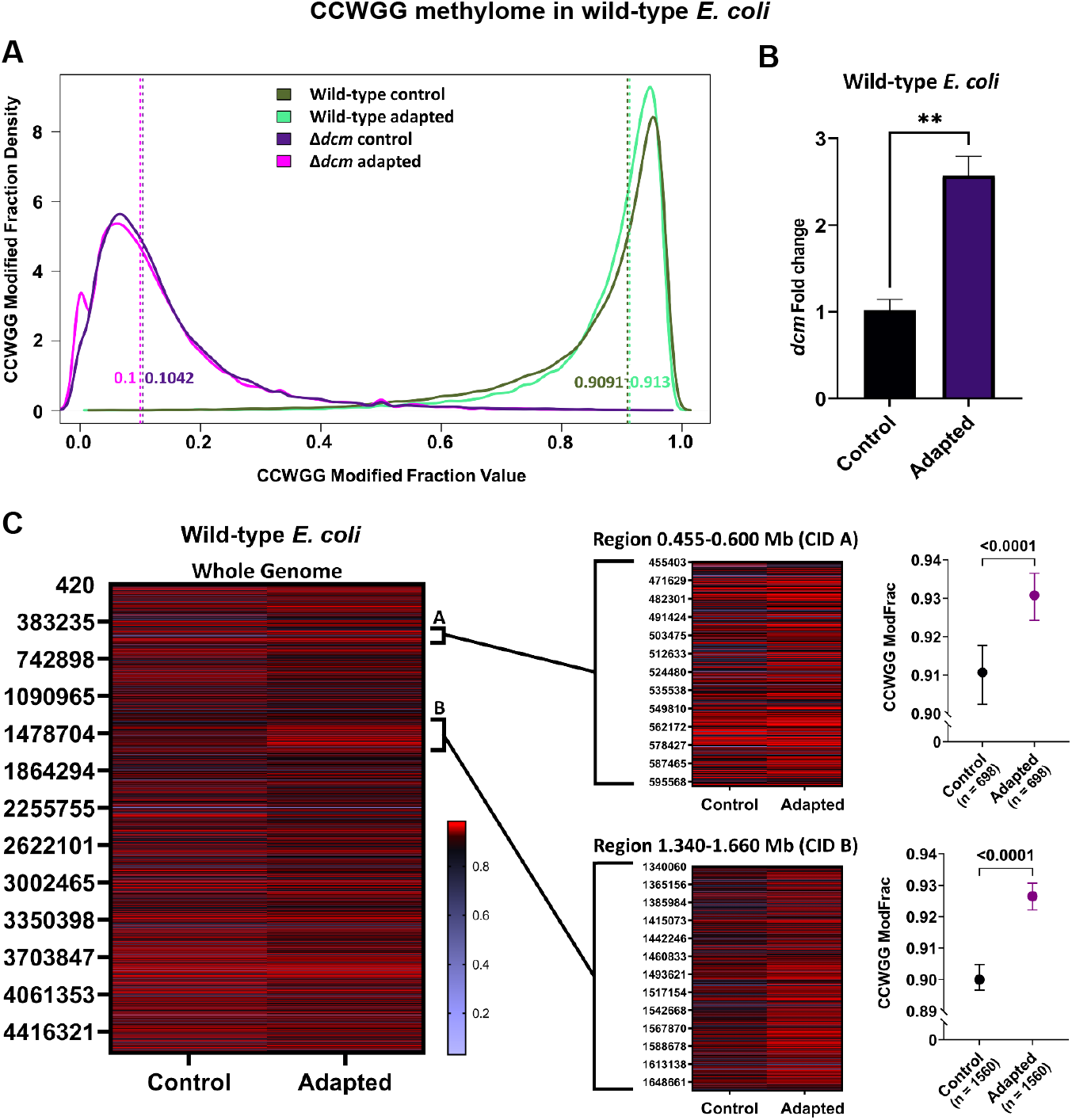
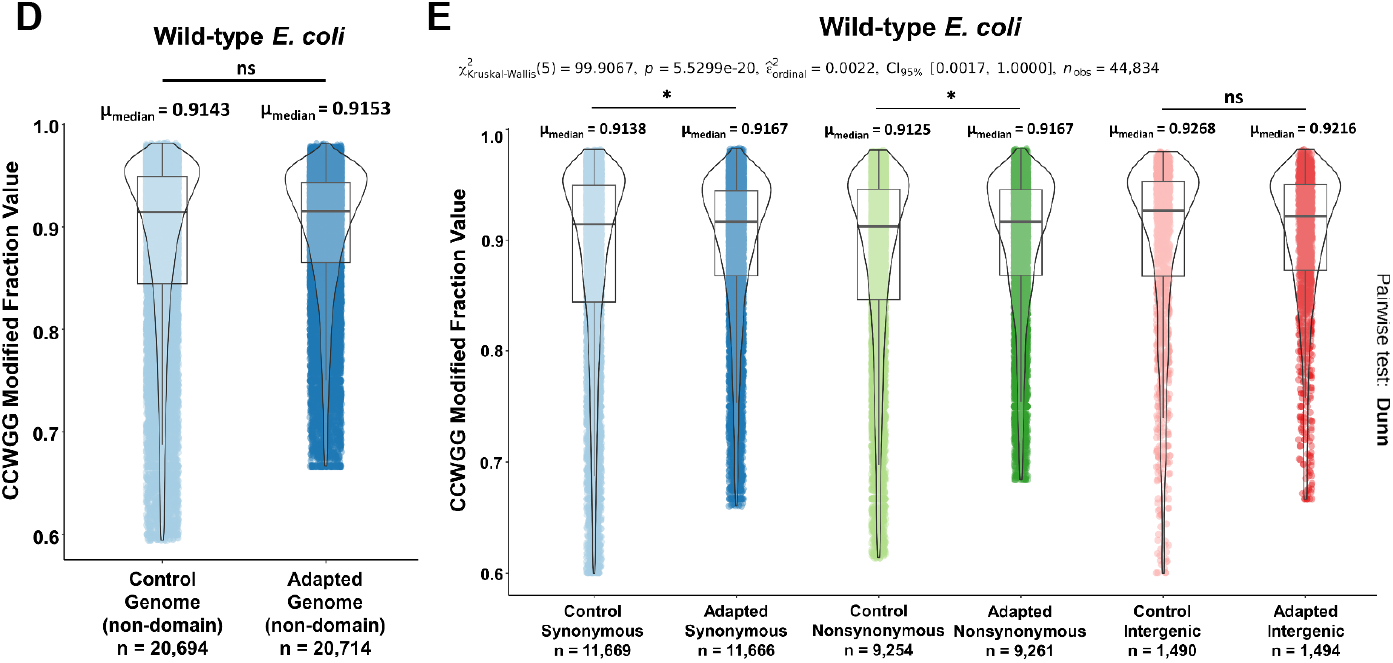
Triclosan-induced changes in the CCWGG (dcm motif) methylome of *E. coli* correlates with chromosome topology changes. **(A)** Kernel density estimation (KDE) plot of modified fraction (ModFrac) values of the internal cytosine in CCWGG motif for wild-type (green) and Δ*dcm* (purple) *E. coli* adapted to triclosan and their respective controls (untreated). The dashed lines denote median ModFrac values. **(B)** *dcm* gene expression is upregulated ∼2.5-fold in wild-type *E. coli* strain in the presence of triclosan stress (two-tailed unpaired Student’s t-test, ** *P*<0.01). Error bars indicate standard error from the mean of three biological replicates. **(C)** Heatmap depicting the ModFrac values for CCWGG sites in adapted and control *E. coli* wild-type strains. The color scale of heatmap corresponds to ModFrac values (maximum=red, median=black, minimum=blue). The right panel shows magnified CID A (0.45-0.60 Mb) and CID B (1.34-1.66 Mb) regions identified from contact maps in Figure 1. Median ModFrac values of the specified regions with 95% confidence interval (error bars) are represented on extreme right. Significantly higher levels of CCWGG methylation can be observed in these regions compared to the control. (Pairwise test: Mann-Whitney). (D) Violin plots showing the distribution of cytosine ModFrac values for genomic regions outside the chromosomal domains (non-domains) observed in triclosan adapted wild-type *E. coli* under triclosan stress and its corresponding control. No significant differences in cytosine methylation were observed (pairwise test: Mann-Whitney; 5^th^ percentile of ModFrac value distribution has been removed from each dataset). **(E)** Triclosan stress induced increase in methylation levels of the internal cytosine in the CCWGG motif in synonymous and non-synonymous sites (position of the internal cytosine) but not in intergenic regions. (5^th^ percentile of ModFrac value distribution has been removed from each of the six datasets). * *P*<0.001.

Since methylated cytosines are prone to spontaneous deamination to thymines and thus act as mutational hotspots (38), we wanted to check if the methylation level of the internal cytosines in CCWGG motif varies based on their position in intergenic regions or within gene bodies. Within the gene bodies, we probed the methylation levels of the internal cytosine in CCWGG motifs at synonymous and non-synonymous sites as these cytosines can be deaminated to thymines. We manually mutated all the CCWGG motifs in the control and adapted wild-type *E. coli* strains and used Snippy for variant calling to classify the type of mutation as synonymous, non-synonymous and intergenic. Cytosine methylation levels for CCWGG motifs present in the intergenic regions was marginally higher in the unadapted control compared to the triclosan-adapted strain, but the difference was not statistically significant (Figure 2E). However, for both synonymous and non-synonymous positions, we observed modest but significant increase in cytosine methylation in the triclosan-adapted strain. While these findings may have important implications, the modest difference in methylation levels at synonymous and non-synonymous sites and the presence of fewer CCWGG motifs in the intergenic regions preclude unambiguous inferences.

In addition to cytosine methylation, we also wanted to interrogate differences in the adenine GATC methylome between control and adapted *E. coli* under triclosan stress. The KDE plot of the ModFrac values for methylated adenine in GATC motifs (Dam methylation sites) in all four strains had peaks at ∼0.9 (Figure 3A). The median modified fraction values are 0.9167, 0.9062, 0.9091 and 0.8889 for wild-type control, wild-type adapted, Δ*dcm* control and Δ*dcm* adapted strains, respectively. We did not find any significant difference in *dam* expression for the adapted wild-type and the adapted Δ*dcm* strains under triclosan stress compared to their respective controls (Figure 3B). Interestingly, when we plotted the heatmaps for all the adapted and control strains, with each row representing the chromosomal position of a GATC motif and its color corresponding to its ModFrac value (Figure 3C & 3D), we found hypermethylation of adenines in the CIDs common to both the adapted strains. Comparing the adenine GATC ModFrac values of these CIDs (CID A & B for wild-type, and CID A & G for Δ*dcm*) with the corresponding regions in controls, we saw significantly higher adenine methylation in the chromosomal domains of the adapted strains (Figure 3C & D summary plots on the extreme right). In contrast, the genomic regions outside the common triclosan-associated domains (CID A&B: wild-type, CID A&G: Δ*dcm*) were associated with significantly reduced adenine methylation levels in both the strains in the presence of triclosan stress (Figure 3E & F). Comparing the distribution of GATC methylation in regions corresponding to the two common CIDs versus the remaining genome (Supplementary Figure S5), it can be observed that the adenine methylation levels are significantly higher in the CIDs for both the wild-type and Δ*dcm E. coli* under triclosan-stress compared to the rest of the genome. As there were more CIDs identified in the Δ*dcm* strain under triclosan stress, which are not found in the wild-type *E. coli*, we assessed the methylation levels of those CIDs as well to compare with the Δ*dcm* control (Supplementary Figure S6). Interestingly, the adenine methylation levels are significantly lower in all the nine domains (nested CIDs have been not assessed separately) which are not common with the wild-type *E. coli*. These results reveal that the adenine GATC methylome changes caused by triclosan stress are quite similar to the cytosine CCWGG methylome differences observed earlier, and both exhibit a relationship with the spatial re-organization of *E. coli* chromosome due to stress induced by triclosan. Our findings highlight the existence of previously unrecognized interplay between spatial re-organization of bacterial chromosomes and the epigenetic landscape under stress conditions.

**Figure 3:**
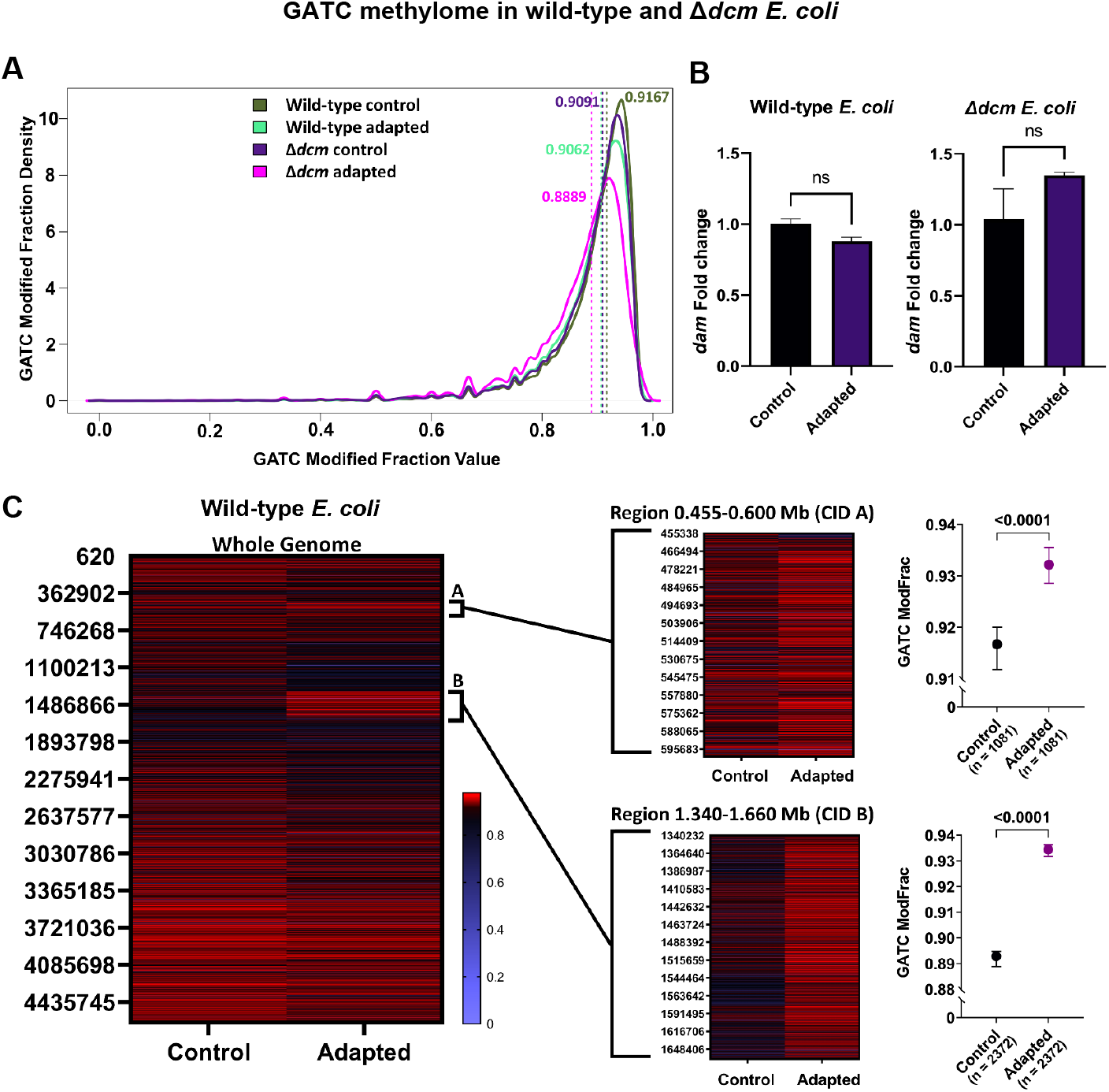
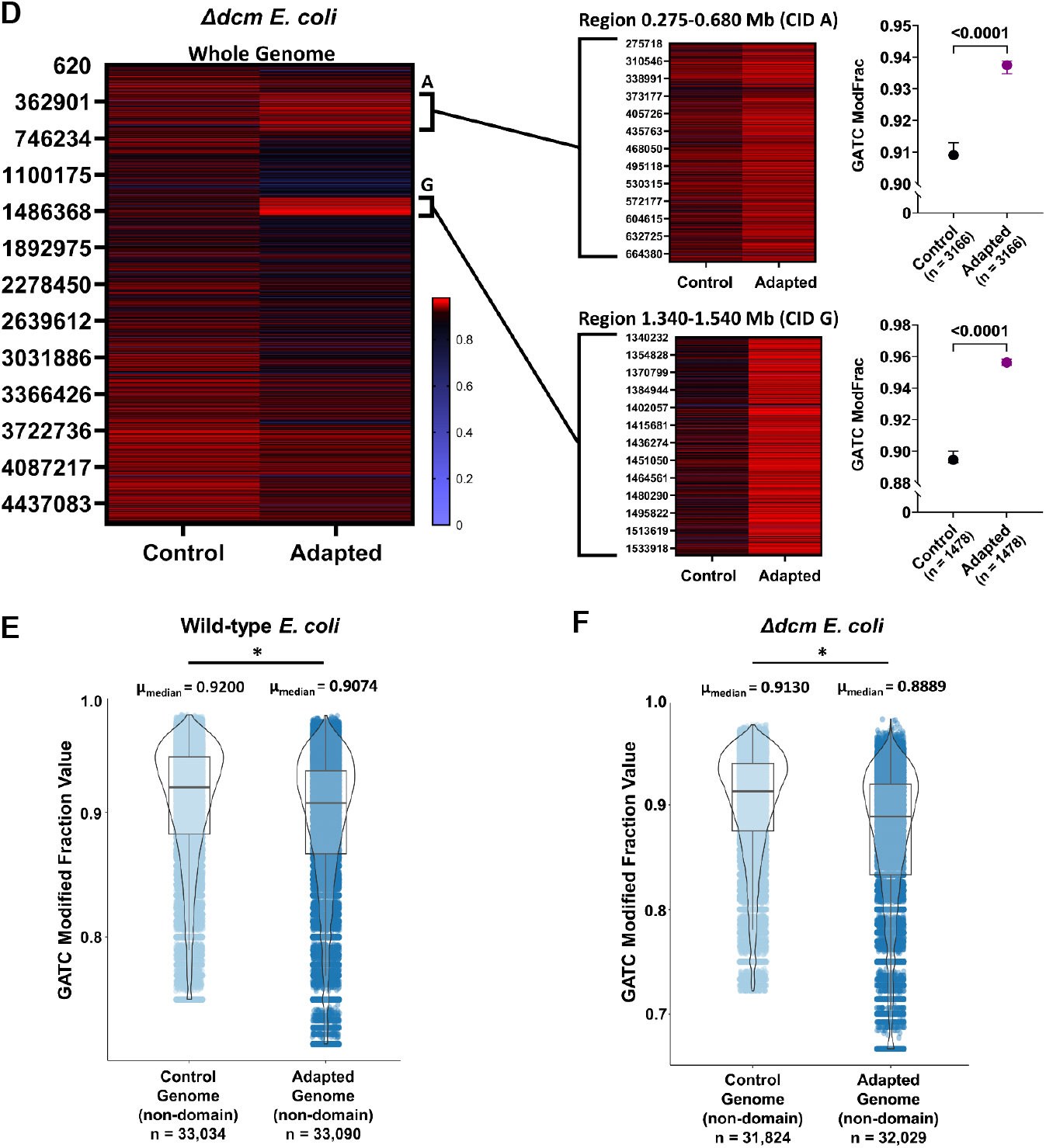
GATC methylome changes in *E. coli* under triclosan stress are linked to changes in chromosome spatial organization. **(A)** Adenine GATC ModFrac values for all adapted and control strains show peaks at ∼0.9 in the KDE plot. Median ModFrac values are denoted by dashed lines. **(B)** No changes in *dam* expression were observed under triclosan stress in both wild-type and Δ*dcm E. coli*. Error bars indicate standard error from the mean of three biological replicates; *P* values are not significant (two-tailed unpaired Student’s t-test). **(C) & (D)** Heatmaps of methylated adenines in GATC motifs in wild-type (C) and Δ*dcm* (D) with y-axes corresponding to their position in the chromosome and colour representing the ModFrac value (maximum=red, median=black, minimum=blue). Smaller heatmaps on the right-side panel are magnified regions of CIDs which are common to both triclosan-adapted wild-type and Δ*dcm E. coli* under triclosan stress. For ModFrac value comparison of the other domains in Δ*dcm E. coli* which do not overlap with the wild-type strain under triclosan stress, please see Supplementary Figure S6. Median ModFrac values with 95% confidence intervals (error bars) on the extreme right show significantly higher levels of adenine methylation within CIDs A & B (wild-type) and CIDs A & G (Δ*dcm*) under triclosan stress. (Pairwise test: Mann-Whitney). **(E) & (F)** Violin plots of GATC ModFrac values from wild-type (E) and Δ*dcm* (F) *E. coli* depicting their distribution in their genomes outside of the common triclosan stress-associated domains (CIDs A & B: wild-type; CIDs A & G: Δ*dcm*). In both the strains, the overall adenine methylation levels outside triclosan stress-associated domains (non-domain) are significantly lower in the adapted strains compared to their controls (pairwise test: Mann-Whitney; 5^th^ percentile of ModFrac value distribution has been removed from each dataset). * *P*<0.0001.

### *E. coli* genes upregulated in triclosan stress are enriched within chromosomal domains associated with triclosan stress

Transcription and higher-order chromosome structures in bacteria are known to be interlinked (11,16,39,40). Previously, it has been shown that under normal growth conditions, increased contact frequency (particularly short-range) strongly correlates with higher transcription levels in *E. coli* (11). To understand how stress-induced changes in chromosome topology correlates with changes in gene expression, we sequenced the total RNA extracted from adapted *E. coli* under triclosan stress and control strains to identify differentially expressed genes (DEGs). By plotting the DEGs along the chromosome (Figure 4A and Supplementary Figure S7.), we found that the transcription profiles of the adapted strains (both wild-type and Δ*dcm E. coli*) under triclosan stress aligned with the two common triclosan-associated chromosomal domains. Of note, genes upregulated in triclosan stress were enriched in the common CIDs (Figure 4B, *left*). We also found that genes down-regulated in triclosan stress were depleted in the common CIDs (Figure 4B, *right*), suggesting that the common CIDs represent chromosomal active genomic regions in *E. coli* during triclosan stress. We also identified the common triclosan-stress associated DEGs in wild-type and Δ*dcm E. coli* (Figure 4C). There were 115 upregulated and 167 downregulated genes common to both adapted wild-type and Δ*dcm* strains under triclosan stress. Plotting these common DEGs along the chromosome (Figure 4D) reflects our previous results showing that upregulated genes co-localize with the triclosan-induced chromosomal domains common to the wild-type and the Δ*dcm E. coli*.

**Figure 4:**
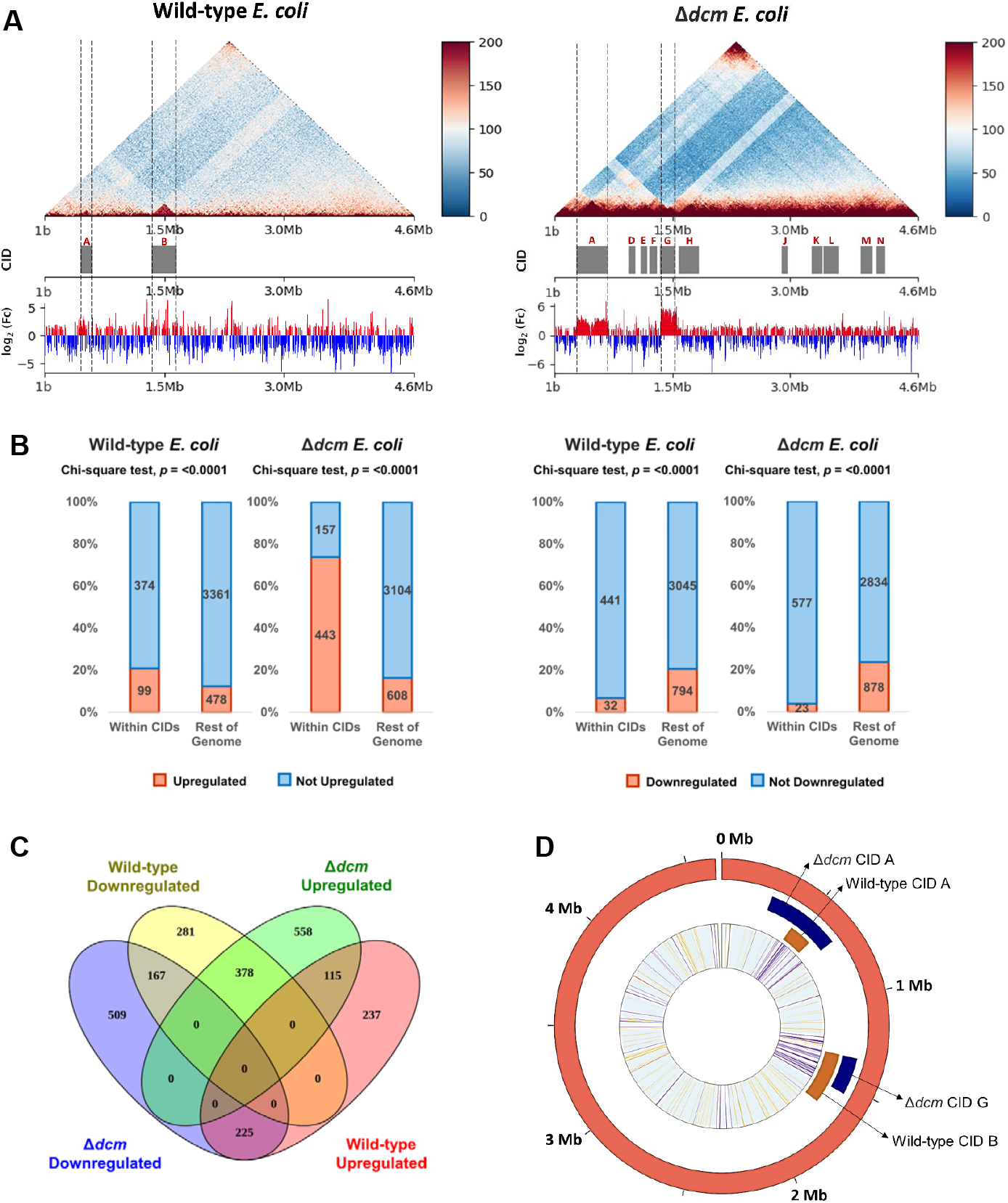
Genes upregulated under triclosan stress are enriched within chromosomal domains, particularly in Δ*dcm E. coli*. **(A)** Interaction matrices annotated with domains (CIDs) and log_2_(Fc = Fold change) values of differentially expressed genes under triclosan stress. The dashed lines indicate the boundaries of common CIDs between wild-type and Δ*dcm E. coli* under triclosan-stress conditions. The color scale of the contact maps corresponds to contact frequency. **(B)** Percent stacked bar plots depicting the number of genes differentially expressed within the CIDs (+/-5 Kb) versus the rest of the genome in wild-type and Δ*dcm E. coli* under triclosan stress compared to their respective controls. Genes upregulated during triclosan stress are significantly enriched within the common triclosan-associated domains compared to the rest of the genome (*left*), while downregulated genes are mostly found outside these common domains (*right*). **(C)** Venn diagram of differentially expressed genes (DEGs) shows there are 115 common upregulated genes and 167 common downregulated genes in wild-type and Δ*dcm E. coli* in the presence of triclosan. **(D)** Circular plot demonstrating the enrichment of upregulated genes in the two common triclosan stress-associated CIDs. The innermost lines represent only common DEGs between wild-type and Δ*dcm E. coli*. Upregulated genes are represented with violet lines, while the yellow lines indicate downregulated genes.

## DISCUSSION

We found triclosan stress induces spatial reorganization of the wild-type as well as Δ*dcm E. coli*. The absence of Dcm methylation (Δ*dcm*) impacts both short-range and long-range chromosomal interactions under triclosan stress. Two of the triclosan-induced CIDs were common to the wild-type and Δ*dcm E. coli*, and one of them had a boundary which overlapped with the *fabI* gene, which encodes the drug target of triclosan. In addition to mutations in the *fabI* gene, triclosan induced other mutations (n=4) and structural variants (n=5). Importantly, all these mutations and structural variants were found within or in close proximity (∼7.5 Kb) to triclosan-induced CIDs. Our findings suggest that triclosan stress induced structural changes (CIDs) are hotspots for genetic variations.

Interestingly, the common CIDs in wild-type and Δ*dcm E. coli* formed under triclosan stress are hypermethylated for both adenine (Dam GA^m^TC motif) as well as cytosines (Dcm CC^m^WGG motif), but similar topology-methylome synergy is absent in *E. coli* growing under normal conditions. DNA methylation in bacteria may either increase or decrease expression of genes depending on the sequence context and gene-specific regulatory mechanisms (6). Here, we find that triclosan stress induced common CIDs, with increased adenine and cytosine methylation, are also transcriptionally active regions enriched in upregulated genes. It is important to note here that these genomic regions which form the triclosan stress-induced CIDs can also harbor large duplications which may explain at least in part, the increased gene expression observed. However, we cannot rule out a possible association between DNA hypermethylation within CIDs and increased gene expression as one of the hypermethylated CIDs in Δ*dcm E. coli* (CID A) did not have any structural variants, but still harbored genes that were upregulated under triclosan stress. The potential role of stress-induced DNA methylation within CIDs in transcriptional regulation among bacteria merits further investigation. We speculate that changes in spatial structures of *E. coli* chromosome under stress conditions allow greater regulatory access to genomic regions which would otherwise have restricted access.

Taken together, our findings identify a previously unrecognized interplay between stress induced chromosome topology changes, generation of genetic variants, epigenetic landscape and transcriptional activity in bacteria. Spatial reorganization of the bacterial chromosome may represent the forefront of the cascade of events that allow adaption to antimicrobial stress.

## MATERIALS AND METHODS

### Strains, growth conditions and adaptation

The wild-type *Escherichia coli* BW25113 (cat# OEC5042) and *dcm* knock-out (cat# OEC4987-200827309) strains used in this study have been obtained from the Keio Knock-out Collection through Horizon Discovery, Cambridge, UK. Glycerol stocks of the *E. coli* cells stored at -80°C were revived by streaking onto Luria–Bertani (LB) agar plates (plates for Δ*dcm E. coli* were also supplemented with 25 µg/ml Kanamycin) and growing overnight at 37°C. Single colonies were inoculated into fresh LB broth and grown at 37°C for subsequent experiments. Minimum inhibitory concentration (MIC) of triclosan in the wild-type and knock-out strains were determined by broth microdilution. Both wild-type and Δ*dcm E. coli* were serially sub-cultured in triplicates with triclosan concentration increasing by 2-fold every second day, starting from a sub-inhibitory concentration of 0.25 µg/ml to 64-fold of their MIC (32 µg/ml). Parallel sub-cultures of both the strains in triclosan-free media were also maintained as controls.

### Growth curve assay

To determine the growth rates of the adapted and control *E. coli* strains, overnight cultures derived from single colonies were diluted 100-fold in fresh media and OD600 values at 37°C were periodically measured using a spectrophotometer. Growth rates were determined using R package growthrates (41) (version 0.8.4; https://CRAN.R-project.org/package=growthrates).

### Antibiotic disc diffusion assay

To test the susceptibility of triclosan-adapted and control *E. coli* strains to different antibiotics, overnight cultures inoculated into fresh LB broth (1:100 dilution) were grown until OD600 = 0.4 and spread onto LB agar plates. Antibiotic discs (listed in Supplementary Table 5) procured from Himedia Laboratories, Mumbai, India were placed on the surface of the inoculated plates. The plates were incubated at 37°C for up to 18-24 hours before measurement of zone of inhibition for each antibiotic.

### Hi-C library preparation and sequencing

*E. coli* Hi-C libraries for single replicates chosen at random were prepared using Proximo Hi-C (Microbe) Kit (Cat # KT1040), Phase Genomics, USA, except the initial cell fixation step. Briefly, to fix the cells, 6 ml of mid-log phase *E. coli* cells, obtained by inoculating fresh media (with 32 µg/ml triclosan for adapted strains) with overnight cultures and growing at 37°C, were collected by centrifugation at 4°C (5000 x g, 10 mins). The pellets were washed with cold 1X PBS and resuspended in 18 ml of 3% (v/v) formaldehyde solution followed by incubation at 4°C for 1 hour on a rocker-shaker. To quench excess formaldehyde, the cells were incubated at 4°C for 15 minutes after adding 2.5 M glycine to a final concentration of 0.375 M. The fixed cells were pelleted by centrifugation, washed with cold 1X TE buffer, flash frozen on dry ice and stored at -80°C until use. For cell lysis and the remainder of the Hi-C library preparation, manufacturer’s protocol was followed. Final libraries were quantified using a Qubit 3.0 Fluorometer and associated Qubit 1X dsDNA HS Assay Kit (Thermo Fisher Scientific, USA). Library quality assessment using an Agilent TapeStation system and paired-end sequencing of the Hi-C libraries were performed at the Quadram Institute Bioscience, Norwich, UK using Illumina NextSeq500 sequencer. The FASTQ files were assessed for quality using FastQC v0.11.9 (https://www.bioinformatics.babraham.ac.uk/projects/fastqc/). Each library had a minimum of 15 million paired reads.

### Generation of contact maps and identification of genomic features

To generate the contact matrix for all the sequenced samples, the paired-end reads from Hi-C sequencing were processed using the CPU version of the automated Juicer (1.6) (23) pipeline. -z, -p and -y flags were used to specify the paths to the reference genome (*E. coli* BW25113; CP009273.1), chromosome size and restriction site files. Since the Hi-C libraries were prepared using the Proximo Hi-C (Microbe) Kit which uses Sau3AI and MluCI restriction enzymes, a custom restriction site file was created using the Python script generate_site_positions.py (https://github.com/aidenlab/juicer/blob/main/misc/generate_site_positions.py). Juicer aligns the reads to the reference and removes duplicate and near-duplicate reads to create contact matrices. Identification of chromosomal interaction domains (CIDs) and loops at 5 Kb, 10kb and 25kb resolutions was done using Arrowhead and HiCCUPS algorithms respectively, which are a part of the Juicer toolkit. The .hic files with MAPQ ≥ 30 generated by Juicer were visualized at 10kb resolution in Juicebox version 1.11.08 (42) after normalization with Knight and Ruiz (KR) matrix balancing method (43). For contact decay analysis, the .hic files with MAPQ ≥ 30 were first converted to the cool format with 5 Kb resolution followed by normalization of all the cool files to the smallest read number of all the matrices using HiCexplorer (44). Python package cooltools (45) was used with the normalized matrices as inputs to generate the contact decay plot.

### Identification of substitutions and indels

The paired FASTQ files from Hi-C sequencing were used as inputs to identify single nucleotide polymorphisms (SNPs) and insertions/deletions (indels) in the control and triclosan-adapted *E. coli* strains. Snippy v4.6.0 (https://github.com/tseeman/snippy) was used with recommended parameters for variant calling with *E. coli* BW25113 (CP009273.1) as the reference genome. The sequencing depth of each sample was calculated using Samtools version 1.16 (46) and BAM files from the Snippy output.

### Native DNA sequencing

Genomic DNA was extracted using QIAamp DNA Mini Kit (Qiagen,Cat. # 51306) using manufacturer’s protocol. Briefly, mid-log phase *E. coli* cells were collected by centrifugation at 5000 x g for 10 mins, followed by lysis and treatment with Proteinase K and RNase A. Spin columns were used to wash and elute gDNA from the lysate. The gDNA was quantified using Qubit 2.0 Fluorometer and associated Qubit dsDNA BR Assay Kit (Thermo Fisher Scientific, USA). The quality was assessed by horizonal agarose gel electrophoresis and ImplenNanoPhotometer N60 (Implen, Germany). The native DNA sequencing library was constructed with the PCR-free Native Barcode Expansion kit (EXP-NBD104) and Ligation Sequencing kit (SQK-LSK109) from Oxford Nanopore Technologies (ONT), UK, as recommended by the manufacturer. In short, 1µg of gDNA per sample was end-prepped using Next FFPE DNA Repair Mix and NEB Next Ultra II End repair/dA-tailing Module from New England Biolabs and each sample was ligated with unique barcodes provided by ONT. Equimolar quantities of the barcoded DNA were pooled followed by adapter ligation and clean-up. The washing steps in-between the protocol were done using AmPure beads (Beckman Coulter, USA). The prepared library was loaded onto an R9.4.1 flow cell (FLO-MIN106D) and sequenced on a MinION (ONT).

### Identification of structural variants

Guppy (version 6.1.7) (ONT) was used for basecalling the raw FAST5 files from native DNA sequencing using high accuracy (HAC) model (dna_r9.4.1_450bps_hac.cfg) as well as adapter trimming and demultiplexing to obtain FASTQ files, which were then used to perform quality assessment in Nanoplot (47). The single-end long reads in FASTQ files with Qscore ≥10 were used for mapping to the reference genome (*E. coli* BW25113; CP009273.1) in Minimap2 version 2.17-r941 (48). The mapped BAM files were sorted and indexed using Samtools version 1.16 (49). Structural variant calling was performed using Sniffles v2_2.0.7(50) followed by filtering of the variants using bcftools (46) view with parameter -i ‘AF>0.3 && SVLEN<1000000 && SVLEN>-1000000’. The circular plots annotated with loops, CIDs, nucleotide variants and structural variants were made using the R package Biocircos 0.3.4 (https://cran.r-project.org/web/packages/BioCircos/) (51). Mobile genetic elements (MGEs) were predicted with https://cge.food.dtu.dk/services/MobileElementFinder/.

### Methylation analysis

The multi-read FAST5 files were demultiplexed and converted to single-read FAST5 files using ont_fast5_api (https://github.com/nanoporetech/ont_fast5_api). Tombo version 1.5.1 (36) was used to annotate the raw files with basecalls followed by mapping the raw signals and associated basecalls to reference genome (*E. coli* BW25113; CP009273.1) using the re-squiggle algorithm. Motif-specific modified base detection was carried out using the “alternative_model” with --alternate-bases dam dcm option, which produced statistics files for both Dam and Dcm motifs. Genome browser compatible text outputs with modified fraction values and coverage for each motif position in the genome were generated from these statistics files using the command “tombo text_output browser_files”. Dampened modified fraction (ModFrac) values for both Dam and Dcm motifs were used for further analysis. Distribution of ModFrac values including Kernel density estimation (KDE) and comparison of ModFrac values within CIDs and remaining genome was analyzed in R using packages ggplot2 (https://ggplot2.tidyverse.org/) and ggstatsplot (52). For investigation of cytosine ModFrac values for positions present in gene bodies and non-coding regions, the cytosines were manually mutated to thymine on a text browser and Snippy v4.6.0 (https://github.com/tseemann/snippy) was used to identify variants. We mapped the position of these variants to the CCWGG positions obtained from Tombo followed by analyzing the distribution with ggstatsplot. Scatter plots for median values with 95% confidence interval and double-gradient heatmaps were made using GraphPad Prism 9.0.

### RNA preparation

Total RNA extraction from *E. coli* was carried out using Invitrogen TRIzol Reagent (Thermo Fisher Scientific; Cat. # 15596018). In brief, mid-log phase cells were collected by centrifugation at 5000 x g for 10 mins (at 4°C) followed by resuspension of the pellets in TRIzol Reagent and storage at -80°C overnight. Next day, the samples were thawed on ice and the cells were mechanically disrupted using 0.1mm Zirconia/Silica Beads (BioSpec; Cat. No. 11079101z) on a BioSpec Mini-Beadbeater-24. This was followed by phase separation using chloroform, recovery of RNA from aqueous phase with isopropanol, washing with 75% ethanol and subsequent solubilization of the RNA in nuclease-free water according to the recommended procedure from TRIzol Reagent manufacturer. Quantification and quality assessment of the isolated RNA were done with ImplenNanoPhotometer N60 and horizonal agarose gel electrophoresis.

### Reverse-transcription quantitative PCR

1 µg of total RNA was treated with DNase-I (New England Biolabs) followed by cDNA synthesis using iScript cDNA synthesis kit (Bio-rad) as per the manufacturer’s protocol. Real-time quantitative PCR was performed on CFX96TM Real-time PCR detection system (Bio-rad) using iTaq Universal SYBR Green Supermix 2X (Bio-rad) in a 10 μl reaction volume with appropriate primers listed in Supplementary Table 6. Relative expression of boundary genes, *dam* and *dcm* were calculated by using the ΔΔCt method with *gyrA* as the internal control (53). Analysis of differences in mean fold change of three biological replicates was performed in GraphPad Prism 9.0.

### Transcriptome analysis

Library preparation using QIAseq Stranded mRNA Lib Kit UDI-C (96) (Qiagen, Cat. # 180443) and paired-end sequencing on Illumina Hiseq X Ten were done by Biokart India Pvt. Ltd. (Bangalore, India). Each library had a minimum of 12 million paired reads. The FASTQ outputs provided were assessed for quality using FastQC v0.11.9. The read pairs were aligned to reference genome (CP009273.1) using bowtie2 (2.5.0) (54). The mapped BAM files were sorted and indexed using Samtools v.1.16 followed by counting the reads with HTSeq version 2.0.2 (55). Differential gene expression analysis was performed in R using NOISeq version 2.40 (56) package wherein the read counts were normalized using Upper Quartile method (57). pnr = 0.2, nss = 5, v = 0.02, replicates = “no” and lc = 1 options were used while running Noiseq. DEGs were recovered with probability threshold, q = 0.95. Visualization of DEGs along the chromosome was done using R packages Sushi v.1.7.1 (58) and Biocircos. Venn diagram was made in Venny 2.1 web tool (https://bioinfogp.cnb.csic.es/tools/venny/).

## Statistical analysis

All statistical analyses were carried out using ggstatsplot package in R (52) and GraphPad Prism v.9. Results are presented within the figures with details in their respective legends.

## Data and materials availability

All Hi-C, native DNA and RNA sequencing datasets have been made available from Sequence Read Archive (SRA), National Center for Biotechnology Information (NCBI), under BioProject accession number PRJNA949223.

## ACKNOWLEDGMENTS

The authors would like to thank David Baker at Quadram Institute Bioscience (QIB), UK for helping sequence the Hi-C libraries. The authors also want to thank Emma Waters at QIB for sharing valuable inputs for preparation of the Hi-C libraries. D.G. is supported by CSIR-Senior Research Fellowship from Council of Scientific and Industrial Research (CSIR), Government of India. D.G. received funds to carry out the Hi-C experiments at the University of East Anglia from the British Council, United Kingdom and Department of Biotechnology (DBT), Government of India under the Newton-Bhabha Fund PhD Placements Programme (Ref: 654911384).

## AUTHOR CONTRIBUTIONS

D.G., P.V., and B.A.E. conceptualized and designed the project. D.G. performed the experiments. D.G., P.V., and B.A.E. were involved in analysis and visualization of the data. D.G. and P.V. wrote the original draft. All authors revised and edited the manuscript. P.V. and B.A.E. supervised the project.

