## Supplementary Figure for "Spatial reorganization of *Escherichia coli* chromosome contextualizes triclosan stress-related genetic, epigenetic and transcriptome changes"

### SUPPLEMENTARY FIGURES

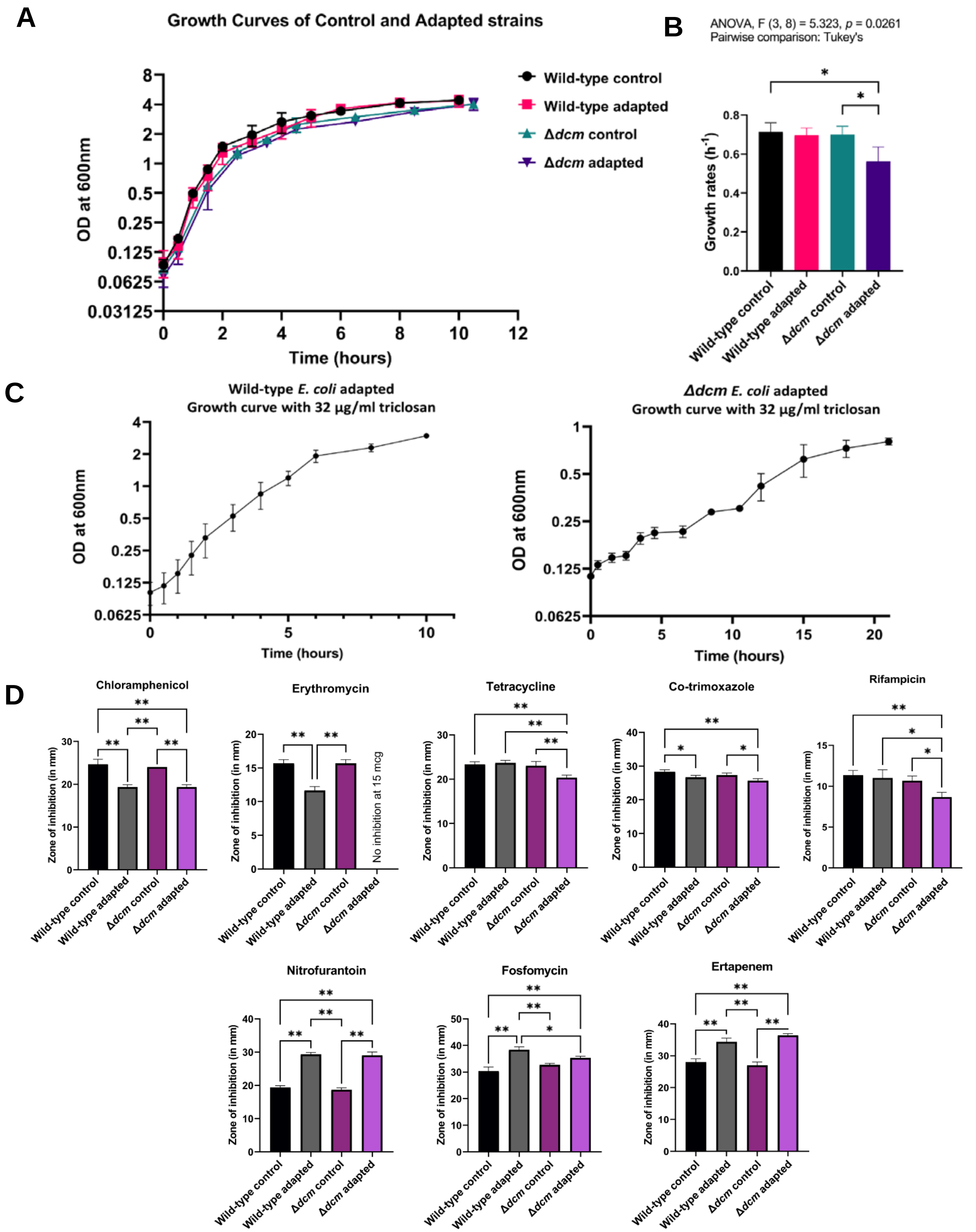


**Supplementary Figure S1.** Phenotypic changes in wild-type and Δ*dcm E. coli* associated with triclosan adaption. **(A)**

The shape of the growth curves for *E. coli* wild-type and Δ*dcm* strains after adaptation to triclosan are comparable to their corresponding controls. This shows that adaptation to triclosan does not alter the entry of *E. coli* into different phases of growth under normal laboratory conditions (n=3; error bars indicate standard error). **(B)** There is a small but significant reduction in the growth rate of *E. coli* Δ*dcm* strain after adaptation to triclosan, but growth rate of *E. coli* wild-type strain is not affected by triclosan adaptation. * *P*<0.05; error bars indicate standard error. **(C)** Growth curves of triclosan-adapted wild-type and Δ*dcm* *E. coli* in the presence of 32μg/ml triclosan. Both the strains have prolonged lag phase and reduced growth in the presence of triclosan stress. **(D)** Altered susceptibility to antibiotics following adaptation to triclosan. *(Top panel)* For *E. coli* wild-type strain, the zone of inhibition is significantly reduced for chloramphenicol, erythromycin and co-trimoxazole after adaptation to triclosan showing reduced susceptibility to these antibiotics. Significant reduction in susceptibility is also observed in triclosan-adapted *E. coli* Δ*dcm* strain for not only chloramphenicol, erythromycin and co-trimoxazole, but also, for tetracycline and rifampicin. *(Bottom panel)* Conversely, significantly increased susceptibility is observed in both the triclosan-adapted wild-type and Δ*dcm* strains to nitrofurantoin, fosfomycin and ertapenem, compared to their corresponding controls. * *P*<0.05, ** *P*<0.01 (Pairwise comparison: Tukey's; n= 3, error bars indicate standard error).

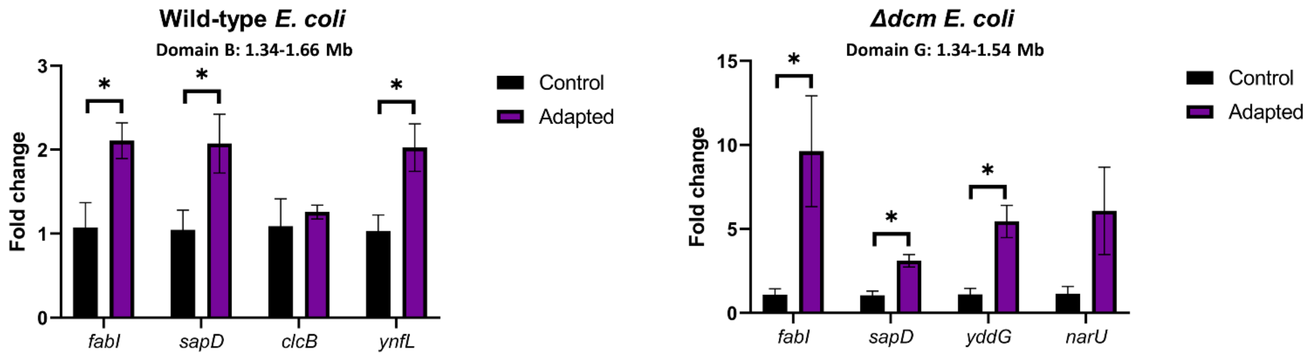


**Supplementary Figure S2.** Expression changes of genes at the boundaries of CID B of wild-type and CID G of Δ*dcm E. coli* in triclosan-adapted strains. *fabI* and *sapD*, which are present at the 1.34 Mb locus, were significantly upregulated in both the triclosan-adapted wild-type and Δ*dcm* strains compared to their corresponding controls. Not all genes at the other boundaries (*clcB* and *ynfL* at 1.66 Mb locus in wild-type, and *yddG* and *narU* at 1.54 Mb locus in Δ*dcm*) were significantly upregulated. (Pairwise test: two-tailed unpaired Student’s t-test; n=3, error bars indicate standard error). * $P<0.05$.

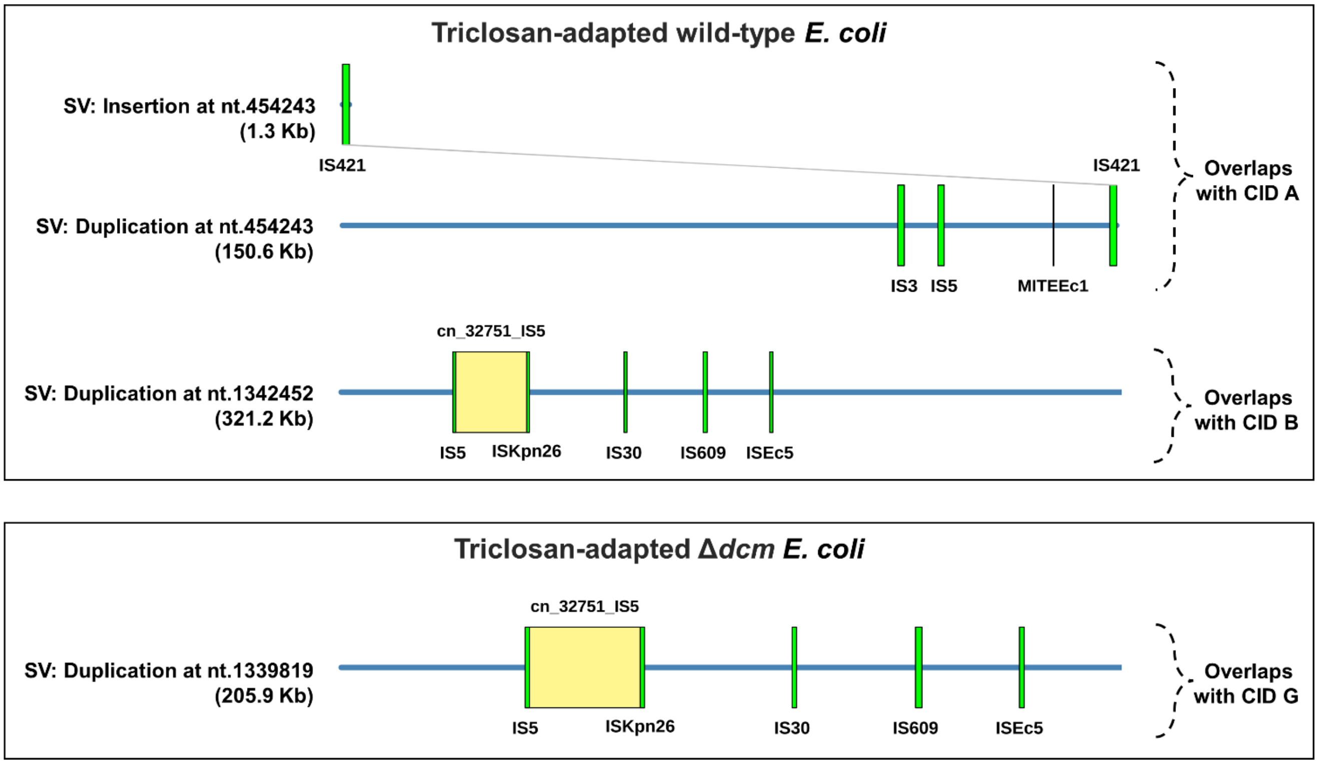


**Supplementary Figure S3.** Predicted Mobile genetic elements (MGEs) present in the structural variants (SVs) identified in triclosan-adapted *E. coli*. In the wild-type strain, the insertion at nt.454243, which is at the starting boundary of CID A, is found to be an Insertion Sequence (IS) element, IS421. This IS element is also found at the end of CID A, at locus 600000 (top panel, gray line indicates matching IS elements). While the large duplications in both strains overlapping CID B (wild-type) and CID G (Δ*dcm*) harbor MGEs, they are not found at the boundaries. All IS elements are indicated in green; composite transposons are indicated in yellow; and the black line represents miniature inverted repeat.

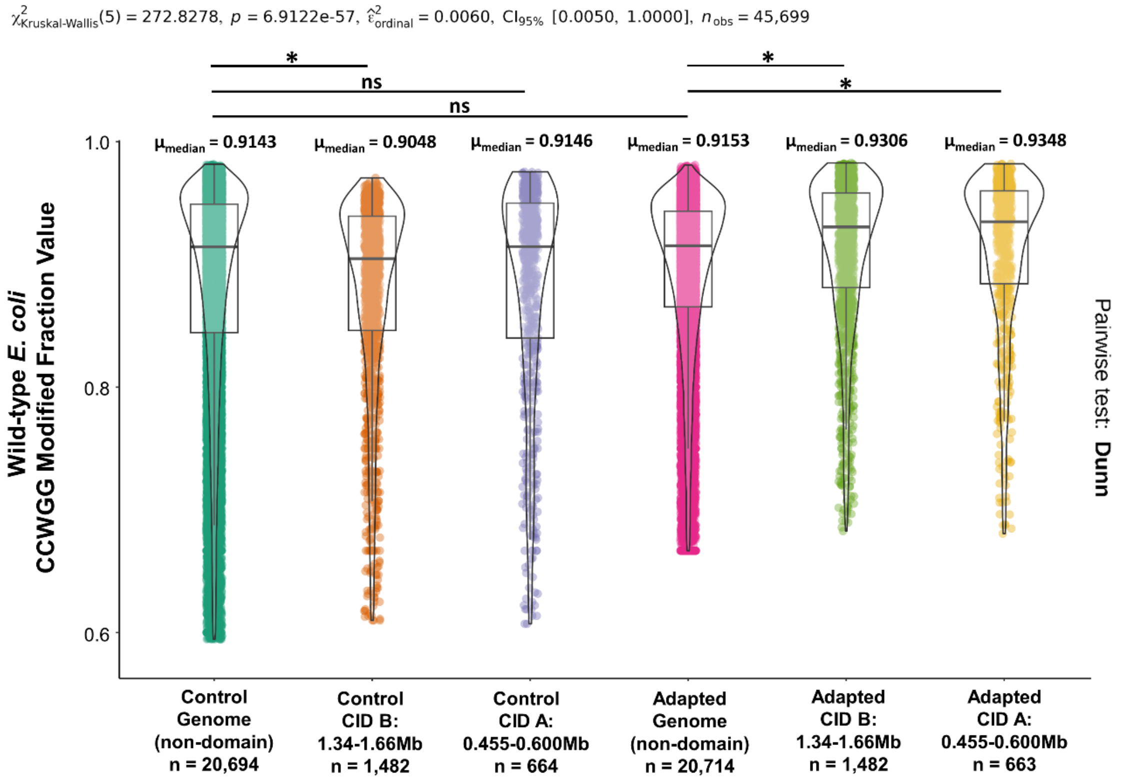


**Supplementary Figure S4.** Violin plots showing the distribution of $CC^{m}WGG$ ModFrac values in CIDs versus rest of the genome observed in triclosan adapted wild-type *E. coli* along with the ModFrac values for the same regions in the unadapted control ($5^{th}$ percentile of the ModFrac value distribution has been removed from each of the six datasets). The median ModFrac values for both CID A and B are significantly higher than the rest of the genome in the triclosan-adapted *E. coli* strain, but the same is not true for the unadapted control. * $P<0.0001$.

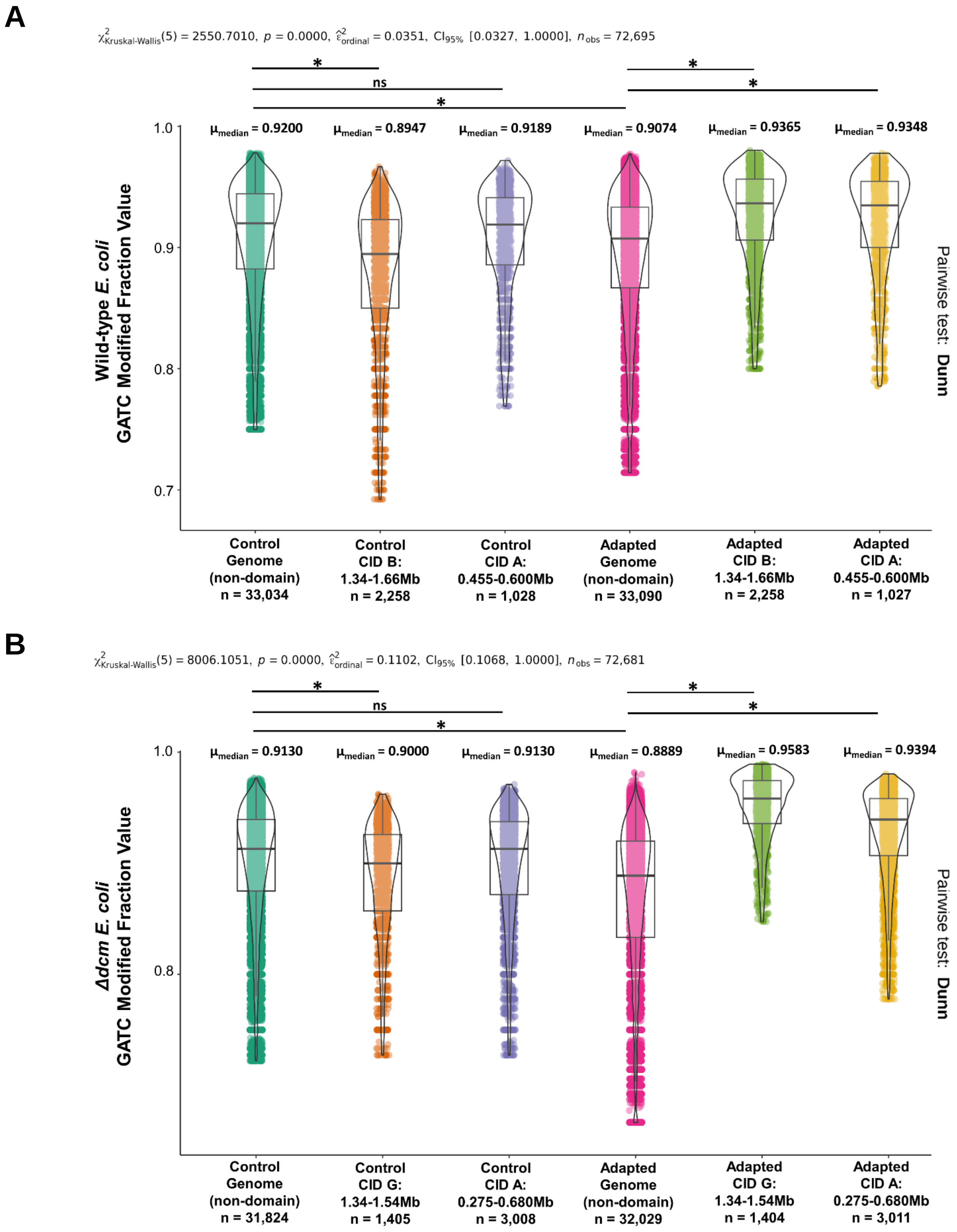


**Supplementary Figure S5.** Violin plots of GA$^{m}$TC ModFrac values from wild-type **(A)** and Δ*dcm* **(B)** *E. coli* depicting their distribution in regions corresponding to triclosan-associated CIDs versus the remaining genome (5$^{th}$ percentile of the ModFrac value distribution has been removed from each dataset). The GA$^{m}$TC ModFrac values are significantly higher in the regions which correspond to CIDs in both the wild-type and Δ*dcm E. coli* under triclosan-stress compared to the rest of the genome. Whereas, in both the controls, no such enrichment of adenine methylation is observed. * $P<0.0001$.

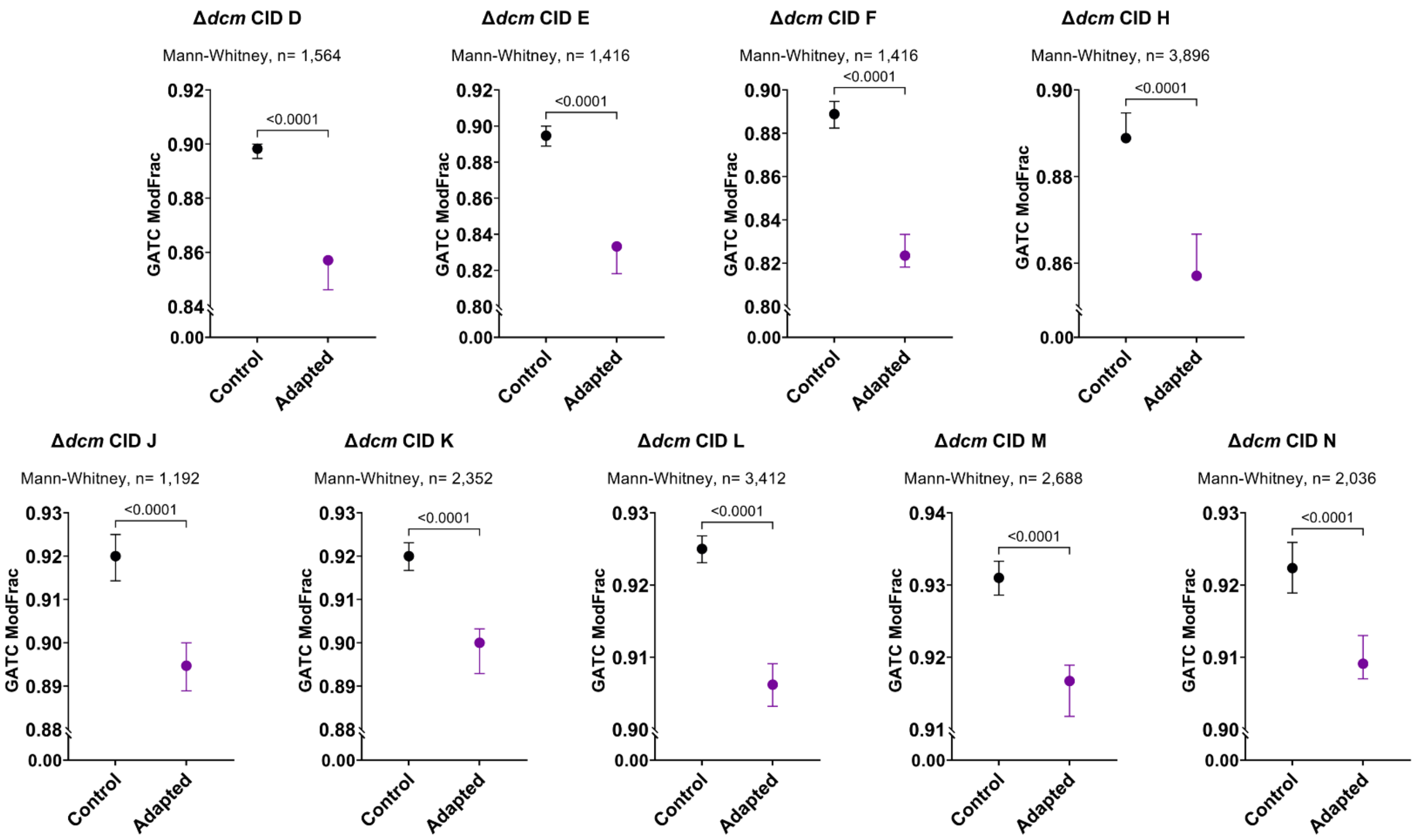


**Supplementary Figure S6.** Comparison of median ModFrac values of $GA^{m}TC$ motifs mapped to chromosomal domains identified in adapted $\Delta$*dcm E. coli* under triclosan stress to the ModFrac values of corresponding regions in the unadapted control. Only CIDs which do not overlap with wild-type *E. coli* under triclosan stress are shown in this figure. In all these domains, the adenine methylation levels are significantly lower in the adapted strain compared to the control, unlike CID A and CID G which have higher adenine methylation levels as seen in Figure 3D. The error bars denote 95% confidence intervals.

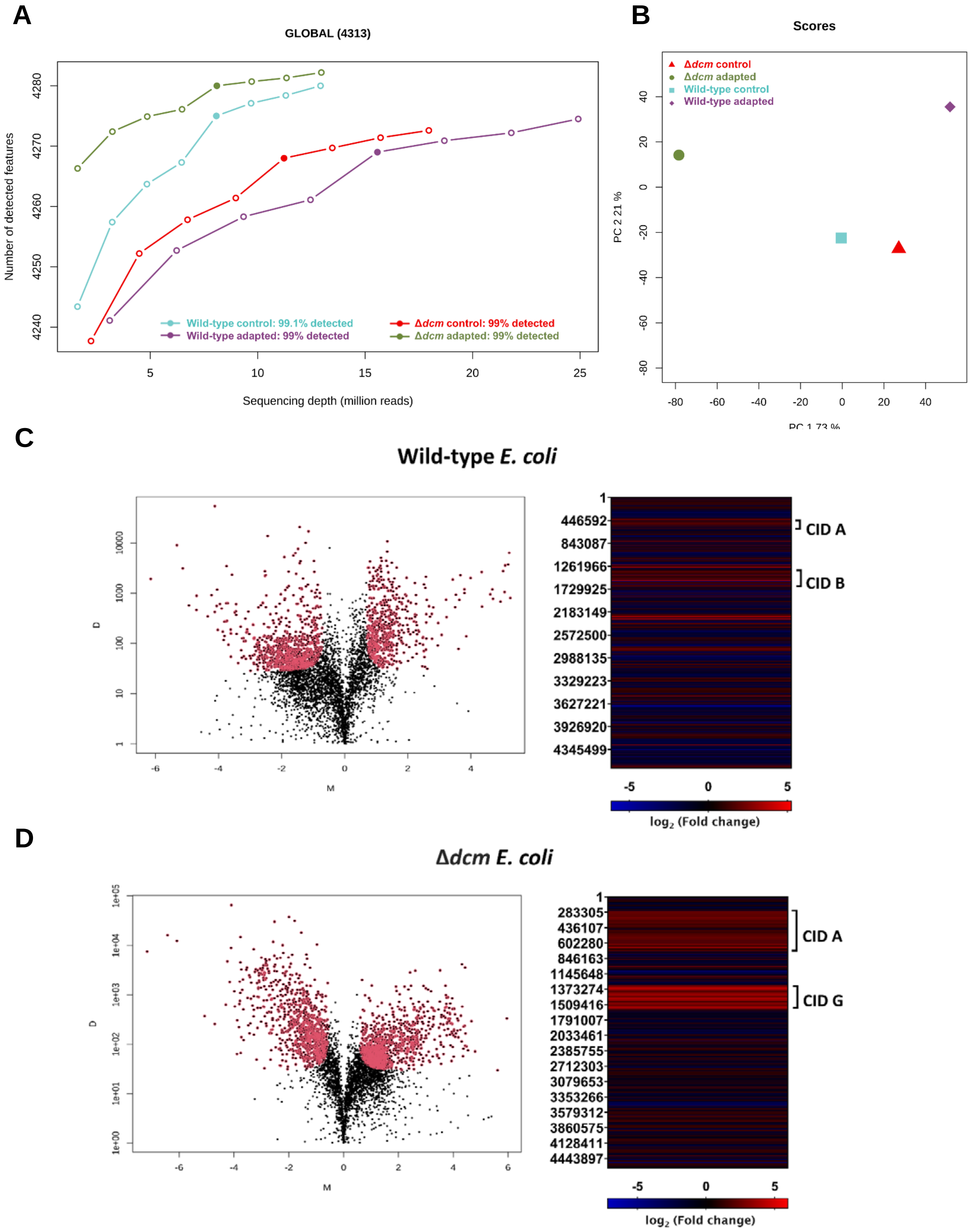


**Supplementary Figure S7.** Quality assessment of RNA-seq read counts and distribution of differentially expressed genes

(DEGs) in *E. coli* under triclosan stress. **(A)** Global saturation plot to compare features detected in all samples with simulated sequencing depths. **(B)** Principal Component Analysis (PCA) showing the controls for both wild-type and Δ*dcm* *E. coli* cluster together indicating similar expression profiles in the absence of triclosan stress. **(C)** & **(D)** The volcano plots depict the $log_2$ fold change (denoted by M on the x-axis) and the absolute difference in expression (denoted by D on y-axis) induced by triclosan stress in wild-type **(C)** and Δ*dcm* **(D)** *E. coli*. DEGs are indicated in red. The probability threshold of differential expression, $q$ = 0.95. The heat maps show the distribution of the DEGs across the genome. The color scale corresponds to $log_2$ fold change values; red indicates upregulated genes, while blue indicates downregulated genes. Common triclosan-associated CIDs have been annotated on the right to show enrichment of upregulated genes within the CIDs in both in wild-type **(C)** and Δ*dcm* **(D)** *E. coli*.

### SUPPLEMENTARY TABLES

**Supplementary Table 1.** Quality assessment of the Hi-C libraries used for preparing the Hi-C contact maps in Juicer.

| Juicer QC Statistics | | | | |
|---|---|---|---|---|
| **Read Pairs** | **Wild-type control** | **Wild-type adapted** | **Δ*dcm* control** | **Δ*dcm* adapted** |
| Sequenced Read Pairs | 26,421,121 | 18,346,398 | 15,190,954 | 21,694,019 |
| Alignable (Normal + Chimeric) | 24,634,833 (93.24%) | 16,933,203 (92.30%) | 13,993,390 (92.12%) | 20,394,178 (94.01%) |
| Unique Reads | 23,121,079 (87.51%) | 14,916,882 (81.31%) | 13,374,486 (88.04%) | 18,285,009 (84.29%) |
| < MAPQ Threshold | 1,040,108 (4.50%) | 1,069,241 (7.17%) | 519,872 (3.89%) | 750,440 (4.10%) |
| Hi-C Contacts | 22,080,971 (95.50%) | 13,847,641 (92.83%) | 12,854,614 (96.11%) | 17,534,569 (95.90%) |
| Short Range (<20Kb) | 4,378,674 (18.94%) | 3,753,902 (25.17%) | 2,820,684 (21.09%) | 6,577,723 (35.97%) |
| Long Range (>20Kb) | 17,694,592 (76.53%) | 10,089,731 (67.64%) | 10,032,579 (75.01%) | 10,954,291 (59.91%) |

**Supplementary Table 2.** Positions of the chromosomal interaction domains (CIDs) identified at 5 Kb, 10 Kb and 25 Kb resolutions from the Hi-C contact maps of each *E. coli* strain **(A)**. Coordinates of loops identified by HiCCUPS in adapted *E. coli* strains under triclosan stress **(B)**.

**A**

| CHROMOSOMAL INTERACTION DOMAINS (CIDS) | | | | |
|---|---|---|---|---|
| **Strain** | **Start locus** | **End locus** | **Arrowhead Score** | **Label** |
| Wild-type control | 420000 | 510000 | 1.74 | A |
| | 2270000 | 2630000 | 1.16 | B |
| | 3250000 | 3435000 | 1.38 | C |
| | 3900000 | 4040000 | 0.81 | D |
| | 4340000 | 4460000 | 1.50 | E |
| Wild-type adapted | 455000 | 600000 | 1.05 | A |
| | 1340000 | 1660000 | 1.71 | B |
| Δ*dcm* control | - | - | - | - |
| Δ*dcm* adapted | 275000 | 680000 | 1.26 | A |
| | 410000 | 510000 | 1.11 | B |
| | 580000 | 675000 | 0.80 | C |
| | 945000 | 1030000 | 0.90 | D |
| | 1105000 | 1185000 | 0.83 | E |
| | 1215000 | 1305000 | 0.74 | F |
| | 1340000 | 1540000 | 1.78 | G |
| | 1585000 | 1845000 | 0.82 | H |
| | 1585000 | 1760000 | 0.82 | I |
| | 2900000 | 2975000 | 0.69 | J |
| | 3280000 | 3410000 | 0.77 | K |
| | 3425000 | 3620000 | 0.78 | L |
| | 3900000 | 4050000 | 0.47 | M |
| | 4095000 | 4215000 | 0.61 | N |

**B**

| Loops identified in adapted strains under stress | | | | |
|---|---|---|---|---|
| **Strain** | **Start locus fragment** | | **End locus fragment** | |
| Wild-type adapted | 100000 | 105000 | 135000 | 140000 |
| | 135000 | 140000 | 170000 | 175000 |
| | 530000 | 535000 | 565000 | 570000 |
| | 1010000 | 1015000 | 1045000 | 1050000 |
| | 1340000 | 1345000 | 1660000 | 1665000 |
| | 2165000 | 2170000 | 2290000 | 2295000 |
| | 2165000 | 2170000 | 2345000 | 2350000 |
| | 2290000 | 2300000 | 2340000 | 2350000 |
| | 2745000 | 2750000 | 2805000 | 2810000 |
| | 3435000 | 3440000 | 3475000 | 3480000 |
| | 3555000 | 3560000 | 3590000 | 3595000 |
| Δ*dcm* adapted | 270000 | 275000 | 680000 | 685000 |
| | 945000 | 950000 | 1025000 | 1030000 |
| | 1340000 | 1345000 | 1540000 | 1545000 |
| | 1520000 | 1530000 | 2980000 | 2990000 |
| | 3665000 | 3670000 | 3710000 | 3715000 |
| | 3910000 | 3915000 | 3960000 | 3965000 |

**Supplementary Table 3. (A)** Substitutions and deletions identified in triclosan-adapted and control *E. coli* wild-type and Δ*dcm* strains, irrespective of association with triclosan adaptation. **(B)** Structural variants identified in control and adapted *E. coli* from long-read nanopore sequencing.

**A**

| Strain (Mean depth) | Position | Gene | Mutation | Fraction of reads | Type |
|---|---|---|---|---|---|
| Wild-type control (563X) | - | - | - | - | - |
| Wild-type adapted (216X) | 1345019 | *fabI* | G>T | 1.0 | Nonsyn |
| Δ*dcm* control (328X) | 190993 | *dxr* | A>T | 0.99 | Nonsyn |
| | 3752869 | *selB* | CGATCGCGTGG | 1.0 | Del |
| | 3940599 | *hdfR* | T | 1.0 | Del |
| Δ*dcm* adapted (582X) | 190993 | *dxr* | A>T | 0.99 | Nonsyn |
| | 3752869 | *selB* | CGATCGCGTGG | 1.0 | Del |
| | 3940599 | *hdfR* | T | 1.0 | Del |
| | 377910 | *insD1* | G>T | 1.0 | Nonsyn |
| | 576681 | *nohD* | G>T | 1.0 | Nonsyn |
| | 1256662 | *prs* | G>A | 0.96 | Nonsyn |
| | 1344690 | *fabI* | T>C | 0.99 | Nonsyn |
| | 4198056 | - | C>A | 0.92 | Intergenic |

**B**

| Strain (Mean depth) | Position | Type | Length | #Reads Reference:Variant | Phred Score (QUAL) |
|---|---|---|---|---|---|
| Wild-type control (135X) | 1203262 | Inversion | 1797 | 73:33 | 59 |
| Wild-type adapted (123X) | 454243 | Insertion | 1342 | 0:88 | 60 |
| | 454243 | Duplication | 150564 | 74:86 | 59 |
| | 1342452 | Duplication | 321245 | 41:113 | 60 |
| Δ*dcm* control (123X) | 1203262 | Inversion | 1797 | 72:38 | 59 |
| | 3174981 | Insertion | 770 | 3:139 | 59 |
| Δ*dcm* adapted (113X) | 1203262 | Inversion | 1797 | 27:57 | 59 |
| | 1339819 | Duplication | 205912 | 0:336 | 59 |
| | 3174981 | Insertion | 774 | 0:75 | 60 |
| | 3627499 | Deletion | 17895 | 0:61 | 59 |

**Supplementary Table 4.** Mobile Genetic Elements (MGEs) predicted in the structural variants (SVs) found in the triclosan-adapted *E. coli* strains.

| Strain | SV | MGE | Position in SV | | Length | Type | Identity | Coverage |
|---|---|---|---|---|---|---|---|---|
| | | | Start | End | | | | |
| Wild-type adapted | INS nt.454243 | IS421 | 13 | 1343 | 1331 | Insertion sequence | 0.930 | 0.963 |
| | DUP nt.454243 | IS421 | 149224 | 150562 | 1339 | Insertion sequence | 0.998 | 0.998 |
| | | IS3 | 107991 | 109248 | 1258 | Insertion sequence | 1 | 1 |
| | | IS5 | 115805 | 116999 | 1195 | Insertion sequence | 1 | 1 |
| | | MITEEc1 | 138196 | 138318 | 123 | Miniature Inverted Repeat | 0.992 | 1 |
| | DUP nt.1342452 | IS609 | 154967 | 156714 | 1748 | Insertion sequence | 0.977 | 1 |
| | | ISEc5 | 183478 | 184768 | 1291 | Insertion sequence | 1 | 1 |
| | | IS30 | 121102 | 122322 | 1221 | Insertion sequence | 1 | 1 |
| | | IS5 | 47850 | 49044 | 1195 | Insertion sequence | 1 | 1 |
| | | ISKpn26 | 79405 | 80600 | 1196 | Insertion sequence | 0.997 | 1 |
| | | cn_32751_IS5 | 47849 | 80600 | 32751 | Composite transposon | 1 | 1 |
| Δ*dcm* adapted | DUP nt.1339819 | IS609 | 157600 | 159347 | 1748 | Insertion sequence | 0.977 | 1 |
| | | ISEc5 | 186111 | 187401 | 1291 | Insertion sequence | 1 | 1 |
| | | IS30 | 123735 | 124955 | 1221 | Insertion sequence | 1 | 1 |
| | | IS5 | 50483 | 51677 | 1195 | Insertion sequence | 1 | 1 |
| | | ISKpn26 | 82038 | 83233 | 1196 | Insertion sequence | 0.997 | 1 |
| | | cn_32751_IS5 | 50482 | 83233 | 32751 | Composite transposon | 1 | 1 |

**Supplementary Table 5.** Antibiotics used for disc diffusion assays.

| Antibiotic | Disc Concentration | Himedia Cat # |
|---|---|---|
| Chloramphenicol | 30 mcg | SD006 |
| Erythromycin | 15 mcg | SD013 |
| Tetracycline | 30 mcg | SD037 |
| Co-trimoxazole | 23.75/1.25 mcg | SD010 |
| Rifampicin | 5 mcg | SD030 |
| Nitrofurantoin | 300 mcg | SD023 |
| Fosfomycin | 200 mcg | SD205 |
| Ertapenem | 10 mcg | SD280 |

**Supplementary Table 6.** Primers used in this study for reverse transcription qPCR.

| Gene | Primer (5'->3') | |
|---|---|---|
| *fabI* | F | GGATCGGACCAGCAGAGATG |
| | R | TATGGGTCTGGCAAAAGCGT |
| *sapD* | F | GGTTTCCACTGTTTGACCGC |
| | R | CTGACGAACCGACCAACTCA |
| *clcB* | F | TGATCGTGGGTCTGCTTTCC |
| | R | CAGCACGGCACACAGTTTAC |
| *ynfL* | F | AACCGCATCCTCTTCAGCAA |
| | R | TATTCTCGGGCTGATGCGAC |
| *yddG* | F | CGCCGACGAGGGCTAATAAT |
| | R | GCCGGGAGTCTGTTATTCGT |
| *narU* | F | ACAAAAATGACCAGCGGTGC |
| | R | TGCTTCGAGCATGGGCAATA |
| *dcm* | F | AAGAAGGCGTGAGTGATGAG |
| | R | CGTCGATAATGCGTACCACA |
| *dam* | F | CGGCGTATCACACAAACAG |
| | R | GCTGCTTATACTGCGTCGA |
| *gyrA* | F | GCCTTCCACGCGTTTTTCTT |
| | R | CCAAAACCGGTCGTGAAACC |
